# multiHIVE: Hierarchical Multimodal Deep Generative Model for Single-cell Multiomics Integration

**DOI:** 10.1101/2025.01.28.635222

**Authors:** Anirudh Nanduri, Musale Krushna Pavan, Kushagra Pandey, Hamim Zafar

## Abstract

**Motivation:** Recently developed single-cell multiomics technologies are enhancing our understanding of cellular heterogeneity by providing multiple views of a biological system. CITE-seq (cellular indexing of transcriptomes and epitopes by sequencing) is one such multiomics assay, with the ability to connect cell states to functions by simultaneously profiling RNA and surface proteins from the same cell. However, the distinct technical characteristics of these data modalities pose significant challenges to their integration into a cohesive representation of cellular identity.

**Results:** Here we present multiHIVE, a hierarchical multimodal deep generative model for inferring cellular embeddings by integrating CITE-seq data modalities. multiHIVE employs hierarchically stacked latent variables as well as modality-specific latent variables to capture shared and private information from the modalities respectively, facilitating integration, denoising and imputation tasks. Extensive benchmarking using gold-standard real and simulated datasets demonstrates multiHIVE’s superiority in integrating CITE-seq datasets. Moreover, multiHIVE outperformed the state-of-the-art methods in imputing missing protein measurements and integration of CITE-seq dataset with unimodal dataset. Using a thymocyte development dataset, we showed that multiHIVE’s cellular embeddings can lead to improved trajectory inference and gene trend identification. Finally, using datasets across development and disease, we demonstrated that factorization of multiHIVE-inferred denoised expression into gene expression programs aids in identifying biological processes at multiple levels of cellular hierarchy.

**Availability and Implementation:** multiHIVE is implemented in Python and is publicly available at https://github.com/Zafar-Lab/multiHIVE.

## Introduction

Single-cell sequencing technologies with their ability to generate high-throughput molecular profiles from thousands of cells have revolutionized our understanding of complex biological systems through detailed characterization of cellular state, heterogeneity, and functions (1; 2). Recently introduced multiomics single-cell sequencing technologies further augment our understanding of cellular identities by enabling the joint profiling of diverse molecular readouts from the same cell (3; 4; 5). Among these multi-omics technologies, CITE-seq (cellular indexing of transcriptomes and epitopes) (3; 6), has enabled the investigation of cellular heterogeneity by simultaneously measuring two distinct modalities: transcriptome (RNA) and surface proteins within the same cells. While gene expression profiles from single cells can characterize cellular heterogeneity (7; 8), surface proteins are more abundant and are directly involved in cell signaling and cell-cell interactions providing complementary insights. The integration of these data modalities has emerged as a critical need in the analysis of such multi-omics datasets for a comprehensive understanding of cellular identities (9).

Recently developed computational methods for the joint analysis of paired multi-omics data rely on matrix factorization (10), nearest-neighbor approaches (11; 12) or variational autoencoders (VAE) (13; 14) for capturing the shared information across modalities. However, integrating different data modalities is challenging due to the distinct complexities inherent to each data type. While RNA expression is high-dimensional and plagued by technical biases, dropouts, and noise, protein abundance is affected by background noise from ambient or nonspecifically bound antibodies. Also data from multiple sources can add unwanted technical variations to the expression profiles. While existing methods try to leverage the shared and complementary information across modalities, matrix factorization methods, due to its linear nature, fails to capture complex biological processes and nearest-neighbor approaches suffer due to poor-quality anchors. While VAE-based methods model the unique noise and technical biases associated with each modality (13), existing methods only focus on capturing a shared latent representation which can result in the suppression of modality-specific information. With the rapid expansion of single-cell multi-omics data generation, there is a pressing need for computational methods that address these challenges efficiently and effectively.

To address these challenges, we present multiHIVE (**multi**-omics integration using **H**ierarchical mult**I**modal **V**ariational auto**E**ncoder), a hierarchical multimodal deep generative variational autoencoder (VAE) model designed to generate robust and interpretable cellular embeddings using both RNA and surface protein modalities. multiHIVE uses a hierarchical variational autoencoder to capture shared information across modalities where the latent variable at each hierarchy encodes information about different biological processes. In addition, multiHIVE also captures modality-specific information using private latent variables for each modality which are learned using non-hierarchical VAE. Using multiple CITE-seq datasets with gold-standard cell type annotations, we demonstrate that multiHIVE outperforms existing methods for the integration of CITE-seq datasets. We further show that multiHIVE can impute missing protein measurements and integrate CITE-seq data with unimodal (e.g., scRNA-seq data) datasets while showcasing superior performance in comparison to existing methods. Using a thymocyte development dataset, we show that cellular embeddings inferred by multiHIVE better captures the trajectory of positively selected T cells. Finally, application of multiHIVE on datasets across development (thymocyte) and disease (breast cancer) further show that the hierarchical latent variables inferred by multiHIVE capture different cell type-specific and across cell type expression programs and thus can further elucidate the cellular identities. Particularly, for the breast cancer dataset, based on the usage of expression programs, we were able to find two subpopulations of cancer associated fibroblasts involved in epithelial to mesenchymal transition and lipid metabolism.

## Methods

### Overview of multiHIVE

multiHIVE is a variational autoencoder-based framework designed to integrate features from two distinct omics modalities into a unified latent space (Figure 1a). multiHIVE employs a novel combination of hierarchical VAE and multimodal VAE for the integration of different omics modalities in the cellular embeddings. Based on the inferred cellular embeddings, multiHIVE can perform several downstream tasks including the integration of multiple CITE-seq datasets, imputation of missing protein measurements, integration of multimodal data with unimodal data (e.g., integration of CITE-seq data with scRNA-seq data), and the inference of cell type-specific and across cell type gene expression programs (Figures 1b-d). In addition, the cellular embeddings can be used for clustering of cells and the inference of celluar trajectory. We start by describing the multiHIVE model and inference method and then we describe the downstream tasks.

**Figure 1.**
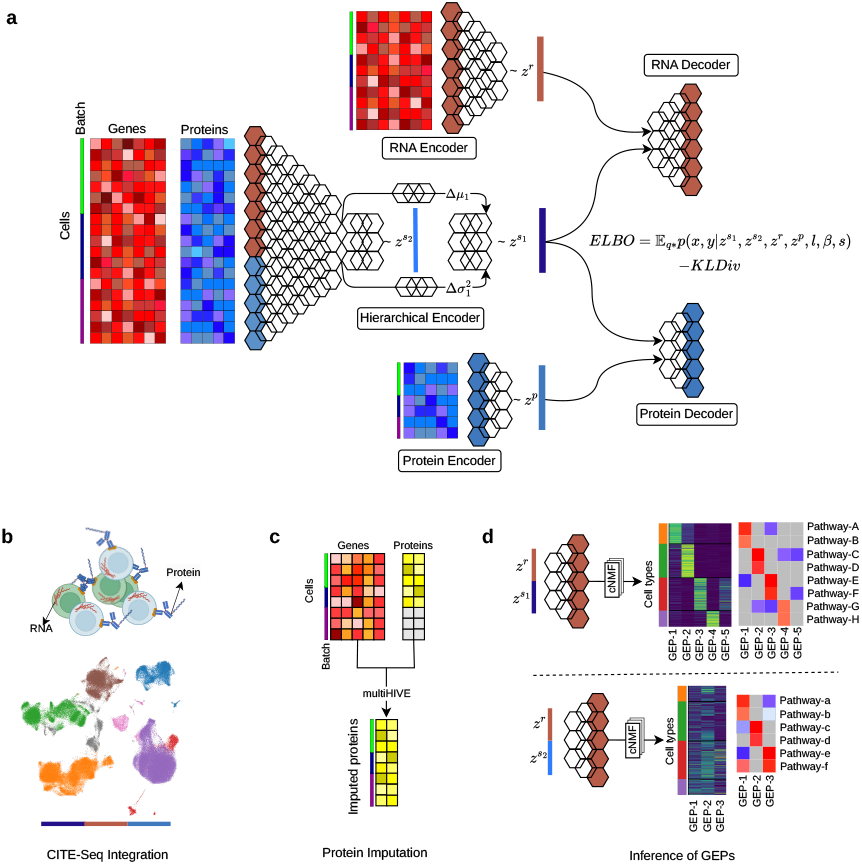
Overview of multiHIVE. (a) The multiHIVE framework consists of a top-down hierarchical variational autoencoder (VAE) for learning a shared latent representation of CITE-seq data, along with two non-hierarchical VAEs for learning private latent embeddings of RNA and protein expression. The model is trained using the Evidence Lower Bound (ELBO) loss function. The hierarchical VAE’s encoder integrates both RNA and protein expression data, while two additional private encoders separately capture RNA and protein-specific latent features. The shared and private embeddings are used to reconstruct RNA and protein expression through two distinct decoders. This combined embedding structure captures the biological relationships between RNA and protein data. The learned embeddings can be applied to several downstream tasks: (b) integration of multiple CITE-seq datasets, along with visualization and clustering, (c) imputation of missing protein measurements, (d) inference of cell type-specific and across-cell type gene expression programs from the RNA reconstructed based on shared latent variables at different hierarchy.

### multiHIVE model

multiHIVE models the joint probability distribution of each cell *n* as ***p***(***x***_*n*_, ***y***_*n*_|*s*_*n*_), where ***x***_*n*_ represents the RNA expression as a *G*-dimensional vector, and ***y***_*n*_ denotes the protein measurements as a *T* -dimensional vector of a cell *n*. The variable *s*_*n*_ contains the batch information (e.g., experimental identifiers) as an one-hot vector. multiHIVE works on a dataset consisting of *N* cells. The estimation of the joint distribution is conducted with a novel variational autoencoder (VAE)-based framework that combines a hierarchical VAE (15; 16) with a multimodal VAE (17). Following the development of multimodal VAEs (17; 18) for joint representation of multi-modal datasets, multiHIVE models the latent representation of the cells using modality-specific latent variables as well as a latent variable shared across all modalities. The modality-specific latent variables for RNA 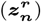, and protein 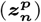 are modeled using non-hierarchical VAEs (19). In addition, multiHIVE incorporates two hierarchical latent variables (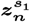and 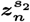) to model the shared information across modalities to effectively capture the complex dependencies in the data as using a hierarchy of latent variables is known to improve the expressive power of VAE (16). Supplementary Figure 1 shows the probabilistic graphical model of multiHIVE.

### Generative Process for RNA

The generative process for RNA for a cell requires the following components: 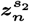, the shared latent representation at level 2; 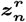, the RNA-specific latent representation; and *s*_*n*_, the batch information. RNA expression for cell *n* and gene *g*, denoted as *x*_*ng*_, follows a negative binomial distribution, which is formulated as a gamma-poisson mixture following previous works (13):

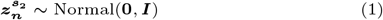

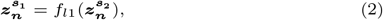

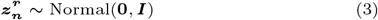

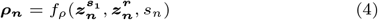

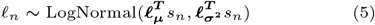

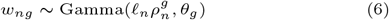

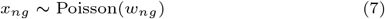

For 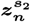 and 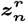, the prior are chosen as isotropic normal distribution. The shared latent representation at level 1, 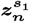 is constructed from 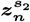 via the neural network *f*_*l*1_. 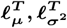 are the mean and variance of library size per batch and these are empirically calculated from the data. The gamma distribution is parameterized by 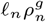 (mean) and *θ*_*g*_ (shape parameter), with 𝓁_*n*_ as a scaling factor. The parameter 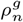 represents the normalized gene expression 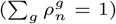. *f*_*ρ*_ denotes the RNA decoder modeled by a neural network. Integrating out *w*_*ng*_ from the above formulation yields the negative binomial distribution for the RNA expression:

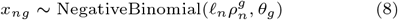

*θ*_*g*_ is interpreted as the inverse dispersion parameter of the negative binomial distribution and also learned in the process of inference (13).

### Generative Process for Protein

Following (13), protein counts are modeled as a combination of background and foreground expression of proteins. The protein count *y*_*nt*_ for cell *n*, and protein *t* is modeled as mixture of negative binomial distributions corresponding to the foreground and background protein expressions. The components of the mixture as well as the mixture weights are reconstructed based on the shared 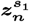 and protein-specific latent representations 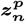 given the batch *s*_*n*_. The generative process for the protein is given as:

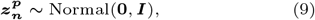

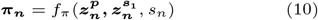

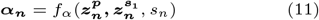

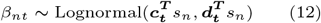

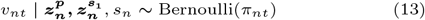

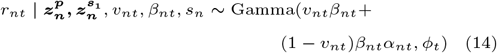

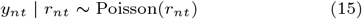

Integrating out *r*_*nt*_ from the above formulation yields

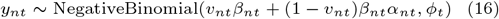

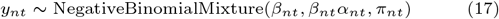

Here, *π*_*nt*_ denotes the probability that the cell–protein pair has observed counts due to background alone and is the output of the neural network *f*_*π*_. *α*_*nt*_ which is the output of the decoder *f*_*α*_ ensures that the foreground is larger than the background. *β*_*nt*_ is the prior probability of the background counts. *v*_*nt*_ controls which mixture component generates the counts considering both background and foreground counts. *ϕ*_*t*_ is the inverse dispersion parameter of the negative binomial distribution for protein *t*. The parameters of the background intensity ***c***_***t***_ and ***d***_***t***_ are learnt during inference and are specific to each protein.

### Hierarchical modeling of the shared latent representation

multiHIVE models the shared information across the modalities using a hierarchy of two latent variables which are modeled using a hierarchical top-down VAE (15; 20). The main idea is to share the top-down path between the variational posteriors and the generative distributions and use a side, deterministic path going from observed variables, as shown in Supplementary Figure 2. According to the schematics in Supplementary Figure 2, the samples from the variational posteriors can be calculated as:

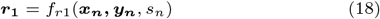

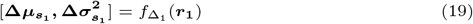

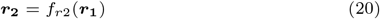

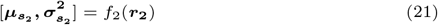

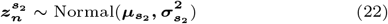

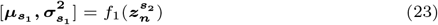

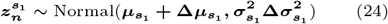

**Figure 2.**
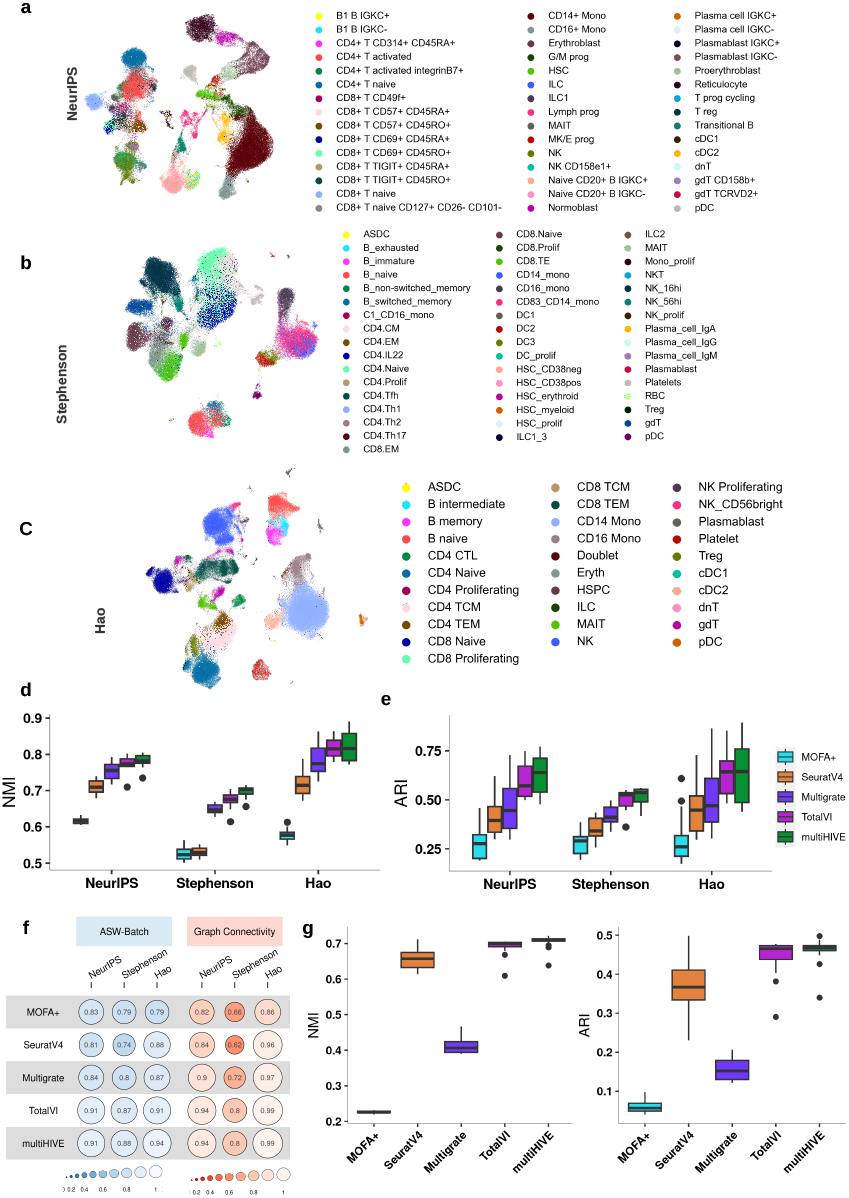
Benchmarking of CITE-Seq integration methods. (a-c) UMAP visualizations of multiHIVE inferred embeddings for three real-world CITE-seq datasets: (a) NeurIPS dataset, (b) Stephenson et al. dataset, and (c) Hao et al. dataset, with cells colored according to their cell type annotations. Comparison of (d) NMI and (e) ARI scores for cell type annotations between MOFA+, SeuratV4, Multigrate, TotalVI, and multiHIVE. For each method, the scores are calculated for 20 different Leiden clustering resolutions ranging from 0.1 to 2.0. (f) Comparison of batch correction metrics ASW-Batch and Graph Connectivity between MOFA+, SeuratV4, Multigrate, TotalVI, and multiHIVE. (g) Comparison of NMI and ARI scores between MOFA+, SeuratV4, Multigrate, TotalVI, and multiHIVE for the simulated dataset simulated using scDesign3 based on the Stephenson et al. dataset.

The parameters in the top-down path are altered by the deterministic bottom-up path and thus helps to encode information of ***x***_***n***_ and ***y***_***n***_ in both the latent variables 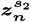 and 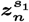. In the above equations, *f*_*r*1_ and *f*_*r*2_ are deterministic functions and *f*_Δ;_1, *f*_1_ and *f*_2_ are variational encoders which are all modeled by neural networks.

### Reconstruction of RNA and protein

The RNA decoder 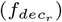 of multiHIVE reconstructs RNA expression of the cells by taking as input the concatenation of the shared representation 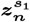 and the RNA-specific latent representation 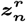, and the batch information

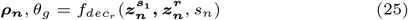

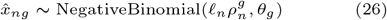

The reconstruction of proteins is done by the protein decoder 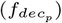 which takes as input the concatenation of the shared representation 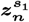 and the protein-specific latent representation 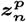

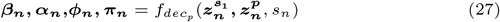

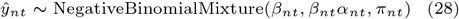

### Inference of latent variables

The posterior distribution of the latent variables is analytically intractable as the model evidence ***p***(***x***_1:*N*_, ***y***_1:*N*_ |*s*_1:*N*_) can not be analytically computed. We use variational inference to approximate the posterior of the model. The variational distribution factorizes as:

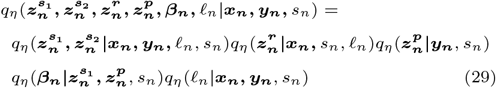

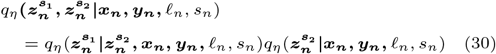

where *η* denotes the set of all parameters of the inference network. The expectations of the approximate posteriors 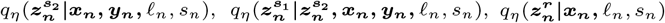, and 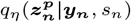 are used as the shared and modality-specific latent cell representations. The model is trained by optimizing the evidence lower bound (ELBO) with respect to variational parameters *η* and model parameters using stochastic gradients (19).

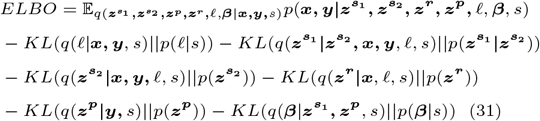

### Training with missing proteins

The missing protein information arises in specific batches *s*_*n*_ either due to the use of different protein panels across batches or when integrating with a unimodal dataset. In both cases, the missing proteins are batch-dependent. Assuming all batches share an identical set of genes, we employed the training procedure described in (21), wherein model parameters are learned exclusively from the observed data. To ensure consistency in data dimensionality at encoders, missing protein values were replaced with zeros, and the optimization of ELBO was based solely on the observed protein values.

The accuracy of imputing missing protein values depends on two factors: (1) the ability of multiHIVE to accurately model the observed protein data, and (2) the statistical distance between the aggregated posterior distributions of 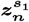 and 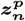 across different batches, as described in (21). A domain adaptation regularization term was added into the ELBO during training, following the approach outlined in (22). This regularization term was scaled by a factor that decayed from 1 to 0 during the early stages of training.

### Downstream tasks

For downstream tasks including clustering, visualization, and trajectory inference, we use a joint embedding, which is obtained by concatenating private and shared latent representations 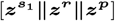.

### Evaluation metrics

To evaluate the performance of integration of CITE-seq dataset, we have considered biological conservation and batch effect correction metrics (23). The biological conservation metrics include normalized mutual information (NMI) and adjusted rand index (ARI) and these utilize the ground truth cell type annotations. The batch correction metrics include average silhouette width-batch (ASW-batch) and graph connectivity and these quantify how well the batches are mixed together.

For the evaluation of protein imputation, we have computed Pearson’s correlation coefficient (PCC) and root-mean square error (RMSE) between the imputed and ground truth protein expression.

### Trajectory inference

We used MARGARET (24) for trajectory inference based on the joint embedding of RNA and protein inferred by multiHIVE. We ran MARGARET with default parameters except for the “obsm data key” parameter which was set to the combined embedding 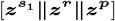 inferred by multiHIVE and “metric clusters” parameter was set to the cell type annotation. The “compute pseudo time()” function was used to infer the pseudotime of cells. To infer the lineage trend of marker genes along the pseudotime, we used the 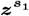 embeddings for reconstructing RNA expression which were then fed to the “plot lineage trends()” function of MARGARET.

### Inference of gene expression programs

To investigate the expression programs captured by each of the lower-dimensional embeddings inferred by multiHIVE, we first trained the multiHIVE model to obtain the lower-dimensional embeddings 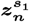 and 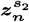. These embeddings are then mapped to the RNA expression space using the trained multiHIVE’s RNA decoder network, producing the reconstructed expression matrices 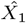 and 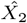 corresponding to 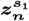 and 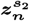, respectively. Consensus non-negative matrix factorization (cNMF) (25) was then used to infer the gene expression programs (GEPs) from the RNA expression matrix reconstructed by multiHIVE. cNMF models the *N ×G* gene expression matrix as a product of two lower-rank matrices, program usage per cell matrix (*N×K*) and a gene expression program matrix (*K ×G*), where *K* is the number of GEPs. The first matrix gives information about the usage of each program in the cell. The second matrix models the contribution of each gene to the program. The value of *K* was determined based on stability and error criteria as suggested by the authors of cNMF (25).

### Pathway analysis

We used fast gene set enrichment analysis (fGSEA) (26) for evaluating the pathways associated with a GEP. The rows of the gene expression program matrix inferred by cNMF gives us a list of genes corresponding to a GEP where the genes are ranked according to their contributions (weights) to the GEP. The ranked list of genes is used as input to fGSEA. In addition, we used the Molecular Signatures Database (MSigDB)(27) to extract the GOBP, REACTOME, and HALLMARK (https://www.gsea-msigdb.org/gsea/msigdb/human/collections.jsp) related pathways (gene sets) and used them as input to fGSEA.

## Results and Discussion

### Benchmarking of multiHIVE for the integration of CITE-seq datasets

To evaluate the performance of multiHIVE in integrating CITE-seq datasets, we applied it on three real-world datasets with ground truth cell type annotation: the NeurIPS dataset (28), the Stephenson et al. peripheral blood mononuclear cell (PBMC) dataset (29) and the PBMC dataset from Hao et al. (11). The NeurIPS Dataset comprises 90,261 single cells derived from the bone marrow of 12 diverse human donors, sampled across multiple sites capturing both developing and differentiated cell types. Stephenson et al. dataset includes PBMCs from a cross-sectional cohort of 130 patients across three UK medical centres. For the purpose of this study, we analyzed the healthy subset of 97,039 cells. The final dataset, Hao et al. is the largest dataset in our analysis, consisting of 161,764 human PBMCs obtained from eight volunteers participating in an HIV vaccine trial. This dataset is generated using a panel of 228 antibodies and profiled using the 10x Chromium platform. In all three datasets, both transcriptomes and antibody-derived tags (ADTs) were profiled from the same cells.

We compared multiHIVE against four state-of-the-art multiomics integration methods: MOFA+ (10), SeuratV4 (11), Multigrate (14), and TotalVI (13). For evaluation, we primarily focused on measuring normalized mutual information (NMI) and adjusted rand index (ARI) for assessing biological conservation based on ground-truth cell type annotation. In addition, we compared the methods based on average silhouette width across batches (ASW-batch) and graph connectivity for evaluating batch correction.

Across all three datasets, multiHIVE was able to capture most of the sub-cell types in different clusters and mix the shared cell types across different batches (Figures 2a-c). Our quantitative comparison showed that multiHIVE consistently achieved the highest NMI value across all datasets at different Leiden clustering resolutions (Figure 2d). multiHIVE was the best-performing method across all datasets based on median

NMI and ARI values as well as the highest NMI and ARI values (Figures 2d,e). In contrast, methods such as MOFA+, SeuratV4 and Multigrate performed poorly in terms of NMI and ARI indicating their inability to preserve the biological cellular identity in CITE-seq integration task. TotalVI was the second-best performing method across all datasets. In terms of batch correction metrics too, multiHIVE and TotalVI outperformed all other methods. For graph connectivity, multiHIVE and TotalVI achieved the same best value across all three datasets, whereas for ASW-batch, multiHIVE outperformed TotalVI on two datasets and performed equally for the NeurIPS dataset (Figure 2f).

For a more robust benchmarking, we further compared the methods on a simulated dataset generated using scDesign3 (30), which learns interpretable parameters from real data. Given the complexity of the Stephenson et al. dataset (indicated by the lower NMI and ARI values of all the methods), we used it as the reference dataset for data simulation using scDesign3. scDesign3 simulated a single-batch synthetic data that closely resembled the real data, incorporating cell-type labels and latent structures. Figure 2g demonstrates that multiHIVE outperformed other methods, achieving the highest NMI and ARI scores on the simulated dataset. TotalVI and SeuratV4 were the second and third-best performer. In contrast, Multigrate and MOFA+ performed poorly indicating limited effectiveness in capturing the underlying data structure.

### Imputation of protein expression by multiHIVE

Given multiHIVE’s ability to integrate different CITE-seq datasets with different protein panels, it can also perform imputation for missing protein measurements. To assess multiHIVE’s protein imputation performance, we used processed PBMCs from two different batches (31; 32; 33). To assess the performance, a subset of proteins was selected as the ground truth, replaced with zeroes, and masked during model training. Then the trained model is used for imputation of the missing values through the protein decoder (Methods). We applied this approach to the PBMC dataset consisting of two batches namely 5k (3,994 cells)(31) and 10k (6,855 cells) (32), comprising 10,849 PBMCs and 14 proteins generated via CITE-seq technology, where the protein counts for one batch were held out during the training and after imputation, the true protein counts were used as ground truth for evaluation. In two different tasks, imputation was performed for the 5k and 10k batches respectively. Imputed proteins were then compared to ground truth protein values using Pearson’s correlation coefficients (PCC) and root mean square error (RMSE) to evaluate the accuracy of the models. We compared multiHIVE’s protein imputation accuracy against that of TotalVI, Multigrate, and SeuratV4 which can also perform protein imputation.

multiHIVE achieved the highest PCC values for 50% of proteins (7 out of 14) for the 5k imputation task (Figure 3a, Supplementary Figure 3a) with values reaching up to 0.98 (e.g., for *CD14*). The second-best performing method, TotalVI, performed best for 5 (out of 14) proteins. SeuratV4’s performance was variable, while it had the highest PCC values for two proteins, it achieved the lowest PCC values for 7 (50%) other proteins. The RMSE scores further highlighted the superior performance of multiHIVE compared to other methods where it had the lowest RMSE values for 8 (out of 14) proteins (Figure 3b).

**Figure 3.**
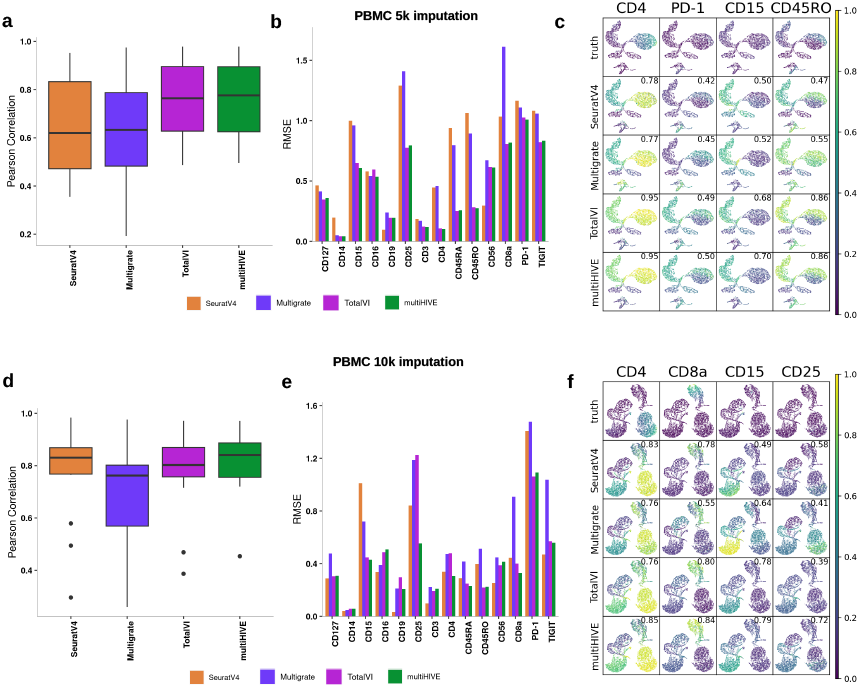
Imputation of missing proteins by multiHIVE. (a-c) Quantitative assessment of imputed protein expression for the 5k batch of the PBMC dataset. Boxplot comparison of (a) Pearson’s correlation coefficient and (b) RMSE values comparing imputed proteins to the ground truth. (c) Feature plots for selected proteins *CD4, PD-1, CD15*, and *CD45RO*. The scatterplot is a UMAP representation of the true protein counts for the 5K PBMC. In each feature plot, the cells in the scatterplot are colored according to the intensity of its relative value for the specified protein. The feature plot color mapping in the first row is based on the true values, whereas in the subsequent rows, it is based on the protein’s predicted expression, as predicted by SeuratV4, Multigrate, TotalVI, and multiHIVE. The number in the top right in each plot is the correlation between the gold standard (true) protein expression counts and the predicted counts. (d-f) Quantitative assessment of imputed protein expression for the 10k batch of the PBMC dataset, with boxplot comparison for (d) Pearson’s correlation coefficient and (e) RMSE. (f) Feature plots for selected proteins *CD4, CD8a, CD15, CD25*. Other descriptions are the same as in (c). In (a)-(f), multiHIVE’s performance is compared against that of SeuratV4, Multigrate and TotalVI.

In the PBMC-10k imputation task, all methods exhibited a general upward shift in correlation values. multiHIVE maintained competitive performance across methods (Figures 3d-f, Supplementary Figure 3b) achieving the highest median PCC. Although Seurat underperformed for the PBMC-5k task, its performance improved for the PBMC-10k imputation. In terms of RMSE also, multiHIVE was the best performer for 5 proteins (Figure 3e) and along with SeuratV4, it outperformed the other methods by achieving lower RMSE. However, for some proteins, SeuratV4 performed very poorly (e.g., *CD15, CD25*, and *PD-1*) (Figure 3e). Figures 3c and 3f visualize the imputed expression for some selected proteins for 5k and 10k imputation which show that multiHIVE’s protein predictions successfully aligned with the underlying cell type embeddings as well as the true protein expression in the data.

### multiHIVE integrates CITE-seq data with scRNA-seq data

Being a generative model, multiHIVE can integrate unimodal scRNA-seq data with CITE-seq data and by using the CITE-seq data as the reference and the scRNA-seq data as the query, multiHIVE enables the prediction of missing protein expression values for the query dataset. To evaluate this, we integrated the human PBMC dataset from (11) with a mucosa associated lymphoid tissue (MALT) dataset generated by 10x Genomics (34) using multiHIVE. Human PBMC dataset consisting of 161,764 cells and 224 proteins served as the reference CITE-seq dataset, whereas the MALT dataset comprising of 8,412 cells and 17 proteins (10 of which overlapped with the proteins profiled in the human PBMC reference) served as the query dataset for which protein expressions were excluded during the training phase. The expression profile of the 10 overlapping proteins were used as ground truth for the evaluation of the accuracy of the protein expression predictions. Initially, the PBMC CITE-seq reference and MALT RNA query data were integrated into a unified latent space, after which the predicted protein expressions were assessed. For this task, we compared multiHIVE against that of SeuratV4 and TotalVI.

The multiHIVE model outperformed other methods in terms of maximum value and median scores for the PCC metric (Figure 4a). Seurat exhibited even negative correlation values for certain proteins. Furthermore, the median RMSE values for multiHIVE was the lowest among the compared methods. Figure 4c visualizes the imputed expression for some selected proteins which shows that multiHIVE’s protein predictions successfully aligned with the underlying cell type embeddings as well as the true protein expression in the data.

**Figure 4.**
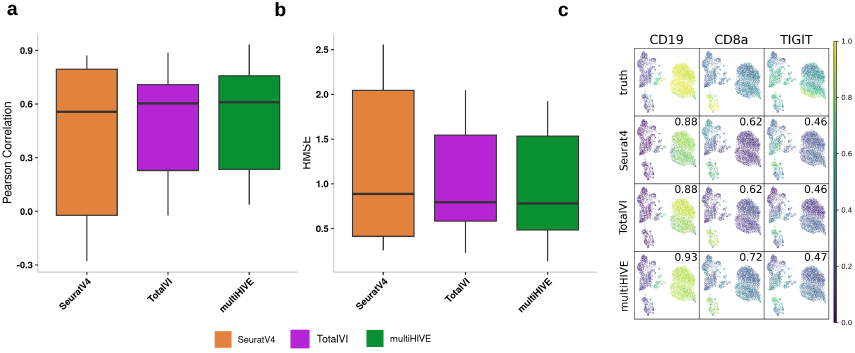
Integration of CITE-seq with scRNA-seq data. (a-c) Quantitative assessment of the integration of human PBMC CITE-seq data (11) with Malt RNA-seq data (34) to predict protein expression levels. Boxplot comparison of (a) Pearson’s correlation coefficient and (b) RMSE values comparing imputed proteins in the Malt dataset to the ground truth. (c) Feature plots for selected proteins *CD19, CD8a*, and *TIGIT*. See Figure 3c for the description of the feature plot. In (a)-(c), multiHIVE’s performance is compared against that of SeuratV4 and TotalVI.

### multiHIVE captures the cellular trajectory and uncovers distinct gene expression programs in thymocyte development

Next, we applied multiHIVE on a thymocyte atlas CITE-seq dataset (35) consisting of 72,042 cells spanning across various developmental stages of T-cells for which whole transcriptome and 111 proteins were profiled. multiHIVE integrated the cells from this dataset and the inferred cellular embeddings captured thymocytes across developmental stages including different double positive (CD4+CD8+) populations (proliferating, quiescent, signaling), immature and mature CD4+ and CD8+ T cells, and CD4-CD8 double-negative cells (Figure 5a). To investigate whether multiHIVE-inferred embeddings captured the continuous developmental process of positively selected thymocytes, we focused on the positively selected thymocytes from DP (Sig.) through mature stages (Figure 5b). Using the multiHIVE-inferred embeddings of these cells and DP (Sig.) cells as the starting population, we performed trajectory inference using MARGARET (24), which inferred a branching trajectory with two different branches leading to mature CD4 and mature CD8 cells as terminal states respectively (Figures 5c,d). We investigated the expression trends of known marker genes over the pseudotemporal trajectory which successfully captured some of the well-known insights, including the early downregulation of *Rag1, Cxcr4*, and *Trbc1*, consistent downregulation of early markers *Ccr9*

**Figure 5.**
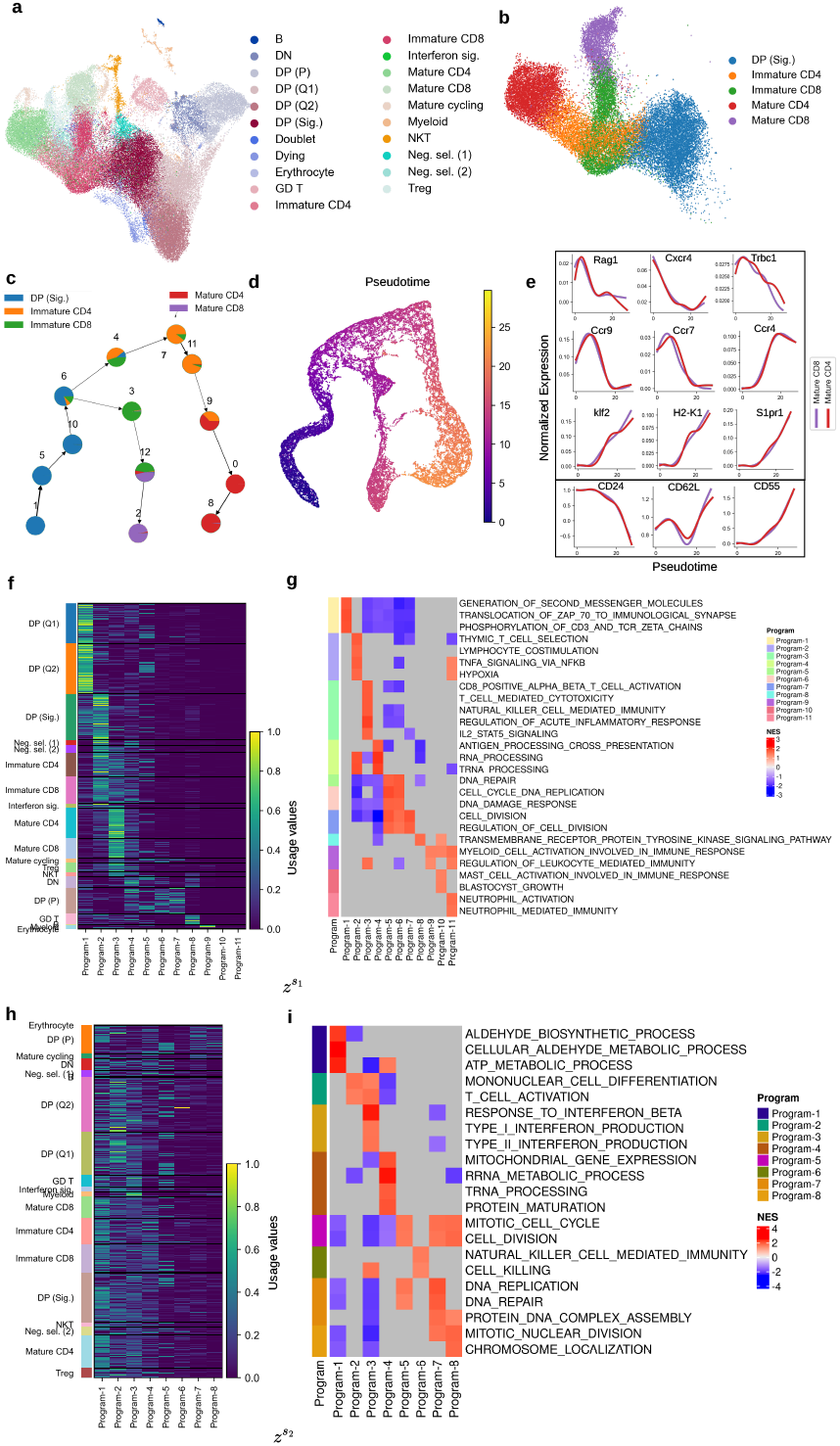
Application of multiHIVE on a thymocyte development dataset. (a) UMAP visualization of multiHIVE embeddings for the Thymocyte development dataset, with cells colored according to cell type. (b) Subset of positive selection cells from (a) for trajectory inference. (c) Trajectory of positively selected CD4+ and CD8+ T cells inferred by MARGARET based on multiHIVE-inferred embeddings. (d) UMAP plot of trajectory delineating the progression of pseudotime. (e) Gene expression trends for known marker genes for the CD4+ and CD8+ T cell lineages using the denoised expression values from MultiHIVE, scaled per gene. (f, h) Heatmaps of the utilization of gene expression programs (GEPs) across all cells, derived using cNMF on RNA expression reconstructed based on multiHIVE embeddings (f) 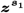 and (f) 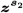 respectively. (g, i) Heatmaps of the normalized enrichment scores (NES) for pathways enriched for the GEPs corresponding to multiHIVE embeddings (g) 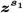 and (i) 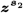 respectively. Gray boxes indicate NA value. and *CD24*, and the subsequent upregulation of maturation markers *Klf2, S1pr1, CD62L* and *CD55* (Figure 5e). We further compared our results to the expression trends over the trajectory reconstructed from cellular embeddings inferred by TotalVI, where the pseudotime order failed to capture the lineage trends of several marker genes (Supplementary Figure 4a-e). For example, *Rag1* exhibited upregulation in the CD8 lineage followed by initial downregulation, *Ccr9* and *CD24* were upregulated after the initial downregulation, and downregulation was observed for the maturation markers *Klf2, S1pr1*, and *CD62L* where it was not expected (Supplementary Figure 4e). Thus, multiHIVE’s embeddings better captured the gene expression changes associated with the branching differentiation of CD4 and CD8 T cells.

We further utilized this dataset to explore the gene expression programs captured by multiHIVE at different hierarchy of the latent space. To do this, multiHIVE-inferred latent representations 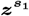 and 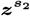 were separately passed through the RNA decoder along with the RNA-specific latent representation ***z***^***r***^ to obtain the reconstructed gene expressions. From the reconstructed RNA expressions, we inferred the gene expression programs (GEPs) using consensus non-negative matrix factorization (cNMF) (25). The pathway enrichment of these GEPs were performed using Fast Gene Set Enrichment Analysis (GSEA) by ranking the genes according to their contributions to the Programs. (26) (Methods).

From the RNA expression reconstructed using 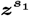 embeddings, we identified 11 GEPs. Despite the complexity of cell type diversity, usage values of these GEPs were found to be cell-type specific (Figure 5f). We observed that Program-1 which had high usage in double-positive (DP) cells (Q1 and Q2), was enriched for pathways relevant to this cell type, such as the translocation of ZAP-70, phosphorylation of CD3 and TCR Zeta chains (Supplementary Table 1). Program-2 had high usage in DP (Sig.) and immature T cells and was associated with pathways such as thymic T cell selection and TNF signaling via NFKB (Supplementary Table 1). Program-3 was mostly used by mature CD4 and CD8 cells and reflected several associated pathways, including CD8-positive alpha-beta T cell activation, T cell-mediated cytotoxicity, and IL2 stat5 signaling (Figure 5d). Similarly, other GEPs were used by different cell types and enriched associated pathways (Program-8 used by gamma-delta T cells enriched for tyrosine kinase signaling, Program-10 and Program-11 used by erythrocytes and myeloid cells enriched for mast cell activation, and myeloid cell activation respectively, see Supplementary Table 1) (Figure 5g).

In contrast, usage values of the GEPs derived from 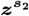 showed a broader distribution across cell types (Figure 5h) suggesting that these GEPs are capturing generic activities across cell types. Pathway analysis of the GEPs further supported this hypothesis, with Program-1 capturing ATP metabolic and aldehyde-related processes, Program-2 capturing mononuclear cell differentiation, Program-3 capturing pathways related to interferon production and responses. Other programs were found to predominantly capture processes related to the cell cycle, cell division, and RNA/DNA-associated pathways (Figure 5i).

### multiHIVE uncovers cell type-specific and across cell type gene expression programs in breast cancer

Finally, we applied multiHIVE on a CITE-seq dataset from Breast Cancer (36) consisting of 7,370 single cells from four experimental batches and whole transcriptome and 118 proteins were profiled for the cells. We first integrated the four batches using multiHIVE which successfully integrated the batches and separated the cell types into different clusters (Figures 6a,b). Applying cNMF on the gene expression reconstructed based on 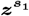 resulted in the identification of 8 GEPs. All these GEPs were cell type-specific as indicated by their usage by different cell types (Figure 6c), for example, program-1 was predominantly used by T cells and B cells, program-2 usage was high for the myeloid cells, program-3 was predominantly used by PVL cells and so on. GSEA analysis of the top genes in these GEPs further enriched pathways related to cell identity (Figure 6d). Program-1 used by T and B cells enriched pathways T cell activation and differentiation. Program-2 was associated with myeloid cell-related pathways such as macrophage activation, myeloid leukocyte activation. Program-4 used by endothelial cells enriched pathways endothelial cell development. Programs 5 and 6 were associated with pathways such as cell matrix adhesion and collagen metabolic processes and were used by cancer associated fibroblasts (CAFs). The last two programs were associated with more general cellular processes, such as cell cycle regulation, and justify their usage value across all cells (Figure 6d). Based on the usage of programs 5 and 6, we further subdivided the CAFs into two subpopulations (CAF P5 and CAF P6) which exhibited higher usage of programs 5 and 6 respectively (Figure 6e, Supplementary Figures 5a-b). Differential gene and protein expression analysis between these two subpopulations revealed that CAF P5 showed elevated expression of collagens (*COL18A1, COL4A1*), *MCAM, HIGD1B, NDUFA4L2* and CCR4 receptor protein (Figure 6f, Supplementsary Figure 5c). In contrast, CAF P6 displayed elevated expression of chemokines (*CXCL1, CXCL14*), growth factors (*IGF1, FGF10*), matrix metalloproteinases (*MMP1, MMP13*), and Nectin-2 transmembrane glycoprotein (Figure 6f, Supplementary Figure 5c). GSEA analysis of these two subpopulations enriched different pathways - CAF P5 enriched lipid metabolism metaprogram and pericyte-like metaprogram (37) and CAF P6 enriched epithelial to mesenchymal transition (EMT) pathway, and hypoxia and CAF-1 and CAF-7 metaprograms (37) (Figure 6g).

**Figure 6.**
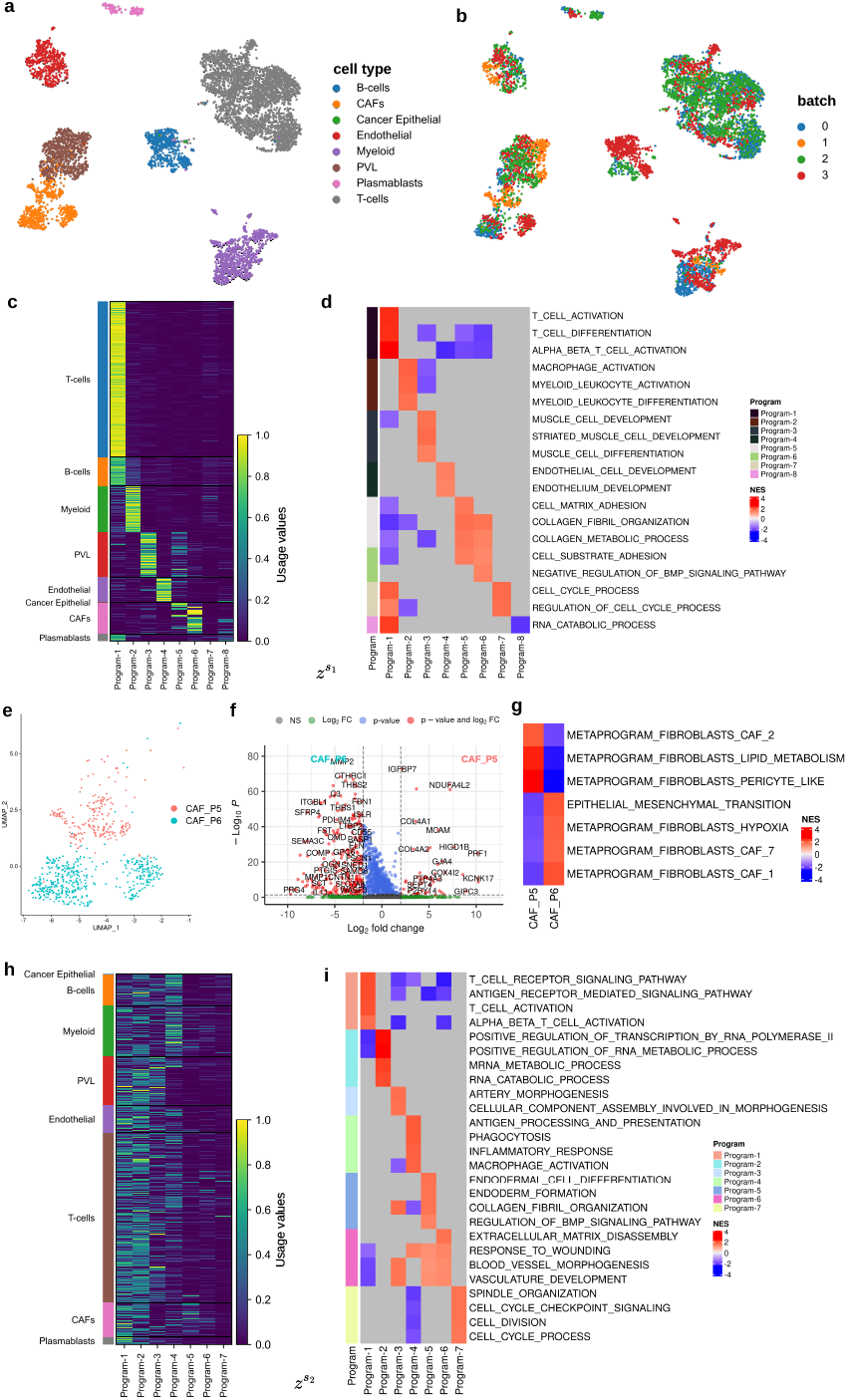
Application of multiHIVE on a breast cancer dataset. (a) UMAP visualization of multiHIVE embeddings for the Breast cancer dataset, with cells colored according to cell type. (b) batch. (c, h) Heatmaps of the utilization of gene expression programs (GEPs) across all cells, derived using cNMF on RNA expression reconstructed based on multiHIVE embeddings (c) 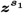 and (h) 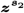 respectively. (d, i) Heatmaps of the normalized enrichment scores (NES) for pathways enriched for the GEPs corresponding to multiHIVE embeddings (d) 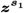 and (i) 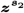 respectively. Gray boxes indicate NA value. (e) UMAP visualization of the CAF subpopulations inferred based on usage of Programs 5 and 6 derived from multiHIVE embeddings 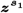 (f) Volcano plot of differentially expressed genes in the CAF subpopulations. (g) Heatmap of the normalized enrichment scores (NES) for pathways enriched for the CAF subpopulations.

In contrast, the 7 GEPs identified from the 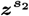 embeddings had usage values broadly distributed across different cell types, indicating that these GEPs represent generic cellular activity programs (Figure 6h). GSEA analysis further uncovered the pathways associated with the GEPs (Figure 6i). While program-1 still enriched pathways related to T cell activation, it also enriched pathways such as antigen receptor mediated signaling. Program-2 was enriched for RNA processing pathways, Program-4 was enriched for phagocytosis and inflammatory response, Program-7 was associated with cell cycle-related pathways. Programs 5 and 6 were used by CAFs and enriched for extracellular matrix disassembly and response to wounding.

## Conclusion

We introduced multiHIVE, a hierarchical multimodal variational autoencoder model for the integration of single-cell CITE-seq datasets. MultiHIVE maps the information from multiple data modalities into a shared hierarchical latent space, capturing cross-modality relationships while also preserving private latent representations specific to each modality.

multiHIVE demonstrated superior performance in integrating CITE-seq datasets compared to state-of-the-art methods MOFA+, SeuratV4, Multigrate, and TotalVI based on both the conservation of cellular identities and the removal of batch effects. multiHIVE is also able to impute missing protein data, enabling integration across datasets with different protein panels. This feature provides valuable insights into proteins that were not detected during experimental procedures. For protein imputation also, multiHIVE achieved better Pearson’s correlation values and lower RMSE scores in comparison with other imputation methods, Multigrate, SeuratV4 and TotalVI. Additionally, multiHIVE can integrate unimodal scRNA-seq datasets with CITE-seq datasets and predict the protein profiles associated with scRNA-seq data. This ability provides a cost-effective alternative to CITE-seq, as RNA-seq experiments are less expensive as compared to multimodal CITE-seq experiments. This feature also opens opportunities for integrating scRNA-seq datasets with inferred protein-level information, enhancing the biological insights gained from such data.

multiHIVE’s unique hierarchical probabilistic model enables the extraction of biological processes at multiple cellular hierarchies. The first-level latent representation captures expression programs associated with cell type, while the second-level representation reveals biological processes occurring across multiple cell types. This was demonstrated using datasets from thymocyte development and breast cancer, where multiHIVE uncovered gene expression programs underlying both cell-type identity (e.g., T cell activation, myeloid leukocyte activation) and cellular activities (e.g. RNA processing, cell-cycle processes). Although single-cell RNA-Seq (scRNA-Seq) can quantify transcripts in individual cells, each cell’s expression profile may be a mixture of both types of programs, making them difficult to disentangle. For the breast cancer dataset, the gene expression programs encoded by multiHIVE’s latent embeddings helped in uncovering two subpopulations of cancer associated fibroblasts which enriched different metaprograms associated to EMT and lipid metabolism (37).

While our primary analysis focused on CITE-seq datasets, multiHIVE’s flexible design allows it to be generalized to other modalities by incorporating modality-specific distributions. This adaptability broadens its potential applications beyond RNA and protein integration. A further future direction would involve the use of hierarchical architecture to explore the multimodal datasets with more than two modalities such as datasets generated using TEA-seq (4) or scNMT-seq (38). With the growing accessibility of single-cell CITE-seq datasets, we anticipate that multiHIVE will empower users to seamlessly integrate and analyze these data, offering a comprehensive view of cells rather than relying on the limited perspective of a single measurement.

## References

1. Stubbington, M. J., Rozenblatt-Rosen, O., Regev, A. & Teichmann, S. A. Single-cell transcriptomics to explore the immune system in health and disease. Science 358, 58–63s (2017).

2. Papalexi, E. & Satija, R. Single-cell RNA sequencing to explore immune cell heterogeneity. Nature Reviews Immunology 18, 35– 45 (2018).

3. Stoeckius, M. et al. Simultaneous epitope and transcriptome measurement in single cells. Nature methods 14, 865–868 (2017).

4. Swanson, E. et al. Simultaneous trimodal single-cell measurement of transcripts, epitopes, and chromatin accessibility using TEA-seq. Elife 10, e63632 (2021).

5. Cao, J. et al. Joint profiling of chromatin accessibility and gene expression in thousands of single cells. Science 361, 1380–1385 (2018).

6. Peterson, V. M. et al. Multiplexed quantification of proteins and transcripts in single cells. Nature biotechnology 35, 936–939 (2017).

7. Regev, A. et al. The human cell atlas. elife 6, e27041 (2017).

8. Shree, A., Pavan, M. K. & Zafar, H. scDREAMER for atlas-level integration of single-cell datasets using deep generative model paired with adversarial classifier. Nature Communications 14, 7781 (2023).

9. Miao, Z., Humphreys, B. D., McMahon, A. P. & Kim, J. Multiomics integration in the age of million single-cell data. Nature Reviews Nephrology 17, 710–724 (2021).

10. Argelaguet, R. et al. MOFA+: a statistical framework for comprehensive integration of multi-modal single-cell data. Genome biology 21, 1–17 (2020).

11. Hao, Y. et al. Integrated analysis of multimodal single-cell data. Cell 184, 3573–3587 (2021).

12. Kim, H. J., Lin, Y., Geddes, T. A., Yang, J. Y. H. & Yang, P. CiteFuse enables multi-modal analysis of CITE-seq data. Bioinformatics 36, 4137–4143 (2020).

13. Gayoso, A. et al. Joint probabilistic modeling of single-cell multiomic data with totalVI. Nature methods 18, 272–282 (2021).

14. Lotfollahi, M., Litinetskaya, A. & Theis, F. J. Multigrate: single-cell multi-omic data integration. BioRxiv 2022–03 (2022).

15. Vahdat, A. & Kautz, J. NVAE: A deep hierarchical variational autoencoder. Advances in neural information processing systems 33, 19667–19679 (2020).

16. Sønderby, C. K., Raiko, T., Maaløe, L., Sønderby, S. K. & Winther, O. Ladder variational autoencoders. Advances in neural information processing systems 29 (2016).

17. Shi, Y., Paige, B., Torr, P. et al. Variational mixture-of-experts autoencoders for multi-modal deep generative models. Advances in neural information processing systems 32 (2019).

18. Wu, M. & Goodman, N. Multimodal generative models for scalable weakly-supervised learning. Advances in neural information processing systems 31 (2018).

19. Kingma, D. P. Auto-encoding variational bayes. arXiv preprint 1312.6114 (2013).

20. Tomczak, J. M. Hierarchical VAEs (2021). URL https://jmtomczak.github.io/blog/9/9_hierarchical_lvm_p1.html.

21. Lopez, R. et al. A joint model of unpaired data from scRNA-seq and spatial transcriptomics for imputing missing gene expression measurements. arXiv preprint 1905.02269 (2019).

22. Ganin, Y. et al. Domain-adversarial training of neural networks. Journal of machine learning research 17, 1–35 (2016).

23. Luecken, M. D. et al. Benchmarking atlas-level data integration in single-cell genomics. Nature methods 19, 41–50 (2022).

24. Pandey, K. & Zafar, H. Inference of cell state transitions and cell fate plasticity from single-cell with MARGARET. Nucleic Acids Research 50, e86–e86 (2022).

25. Kotliar, D. et al. Identifying gene expression programs of cell-type identity and cellular activity with single-cell RNA-Seq. Elife 8, e43803 (2019).

26. Korotkevich, G. et al. Fast gene set enrichment analysis. Biorxiv 060012 (2016).

27. Liberzon, A. et al. Molecular signatures database (MSigDB) 3.0. Bioinformatics 27, 1739–1740 (2011).

28. Luecken, M. D. et al. A sandbox for prediction and integration of DNA, RNA, and proteins in single cells. In Thirty-fifth conference on neural information processing systems datasets and benchmarks track (Round 2) (2021).

29. Stephenson, E. et al. Single-cell multi-omics analysis of the immune response in COVID-19. Nature medicine 27, 904–916 (2021).

30. Song, D. et al. scDesign3 generates realistic in silico data for multimodal single-cell and spatial omics. Nature Biotechnology 42, 247–252 (2024).

31. 10x Genomics. 5k peripheral blood mononuclear cells (pbmcs) with protein expression (2019). URL https://support.10xgenomics.com/single-cell-gene-expression/datasets/3.0.2/5k_pbmc_protein_v3. Accessed via scvi-tools usingscvi.data.pbmcs10xciteseq().

32. 10x Genomics. 10k peripheral blood mononuclear cells (pbmcs) with protein expression (2019). URL https://support.10xgenomics.com/single-cell-gene-expression/datasets/3.0.0/pbmc_10k_protein_v3. Accessed via scvi-tools usingscvi.data.pbmcs10xciteseq().

33. Gayoso, A. et al. A Python library for probabilistic analysis of single-cell omics data. Nature Biotechnology (2022). URL 10.1038/s41587-021-01206-w.

34. (2018). URL https://www.10xgenomics.com/datasets/10-k-cells-from-a-malt-tumor-gene-expression-and-cell-surface-protein-3-standard-3-0-0.

35. Steier, Z. et al. Single-cell multiomic analysis of thymocyte development reveals drivers of CD4+ T cell and CD8+ T cell lineage commitment. Nature Immunology 24, 1579–1590 (2023).

36. Wu, S. Z. et al. A single-cell and spatially resolved atlas of human breast cancers. Nature genetics 53, 1334–1347 (2021).

37. Gavish, A. et al. Hallmarks of transcriptional intratumour heterogeneity across a thousand tumours. Nature 618, 598–606 (2023).

38. Clark, S. J. et al. scNMT-seq enables joint profiling of chromatin accessibility DNA methylation and transcription in single cells. Nature communications 9, 781 (2018).

