## Supplementary Material for "multiHIVE: Hierarchical Multimodal Deep Generative Model for Single-cell Multiomics Integration"

January 23, 2025

#### Contents

**Supplementary Figures** **2**

**Supplementary Tables** **6**

#### Supplementary Figures

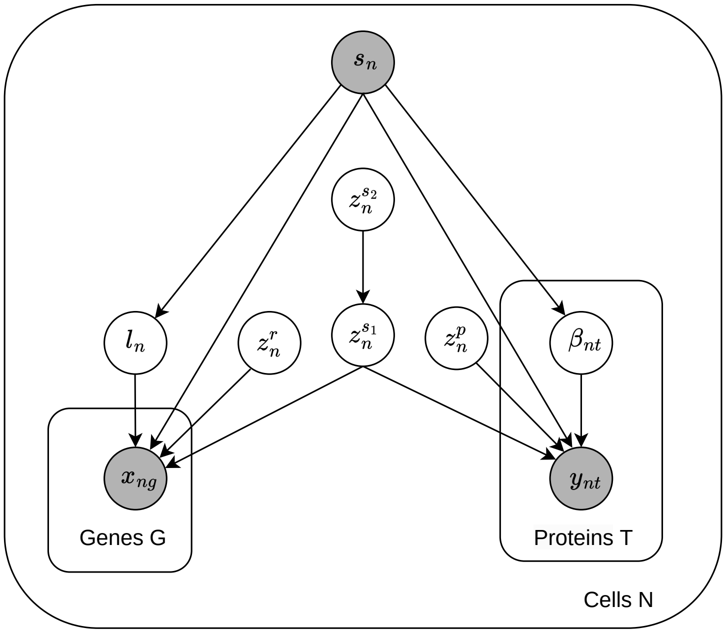


**Supplementary Figure 1**: multiHIVE's probabilistic hierarchical graphical model. Shaded nodes represent observed random variables. Unshaded nodes represent latent variables. Edges denote conditional independence in the direction shown. Rectangles ("plates") represent independent replication of variables inside.

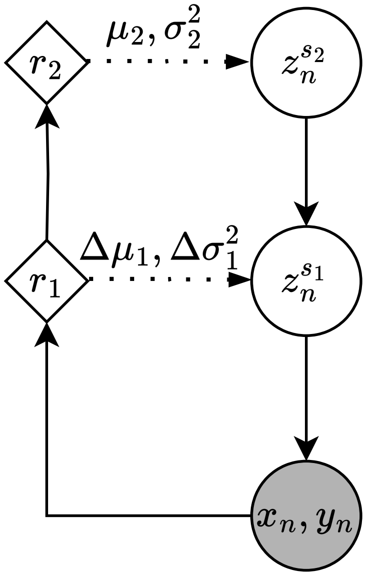


**Supplementary Figure 2**: multiHIVE's architecture for joint latent space representation. The left side of the figure consists of a bottom-up deterministic path $[(x_n, y_n) \rightarrow r_1 \rightarrow r_2]$, and to the right, there is the top-down stochastic path $[z_n^{s_2} \rightarrow z_n^{s_1} \rightarrow (x_n, y_n)]$. The diamond shape indicates neural network transformations. The circle indicates a random variable. The shaded circle indicates observed random variable.

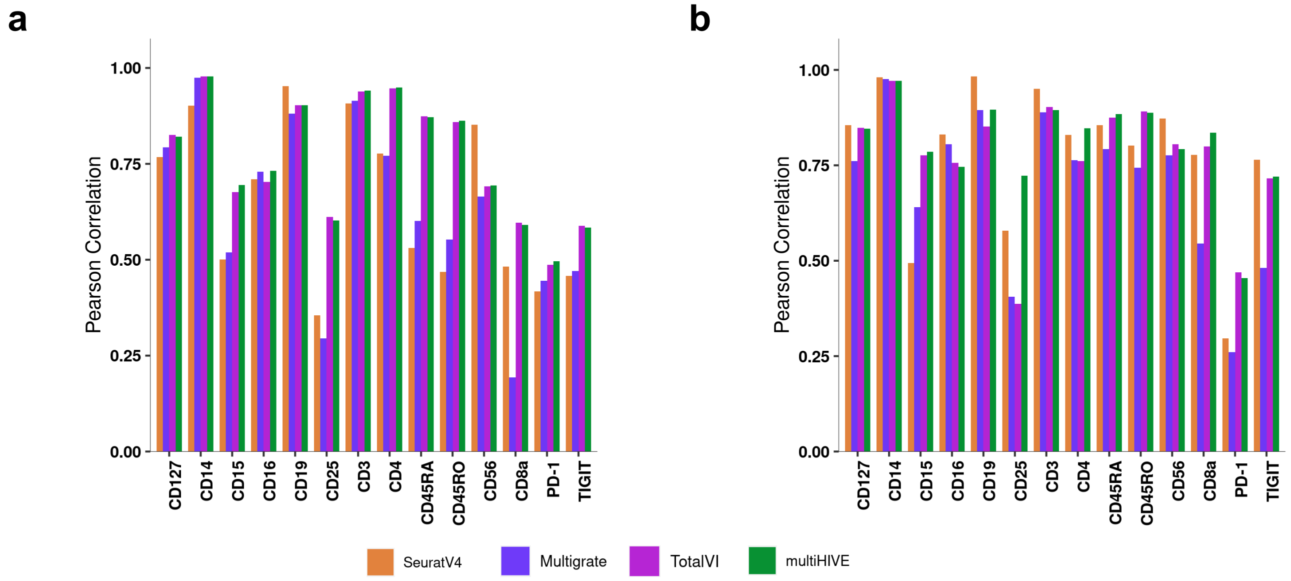


**Supplementary Figure 3**: Barplot comparison of Pearson's correlation coefficient of imputed protein expression for (a) 5k batch and (b) 10k batch of the PBMC dataset. multiHIVE's performance is compared against that of SeuratV4, Multigrate and TotalVI.

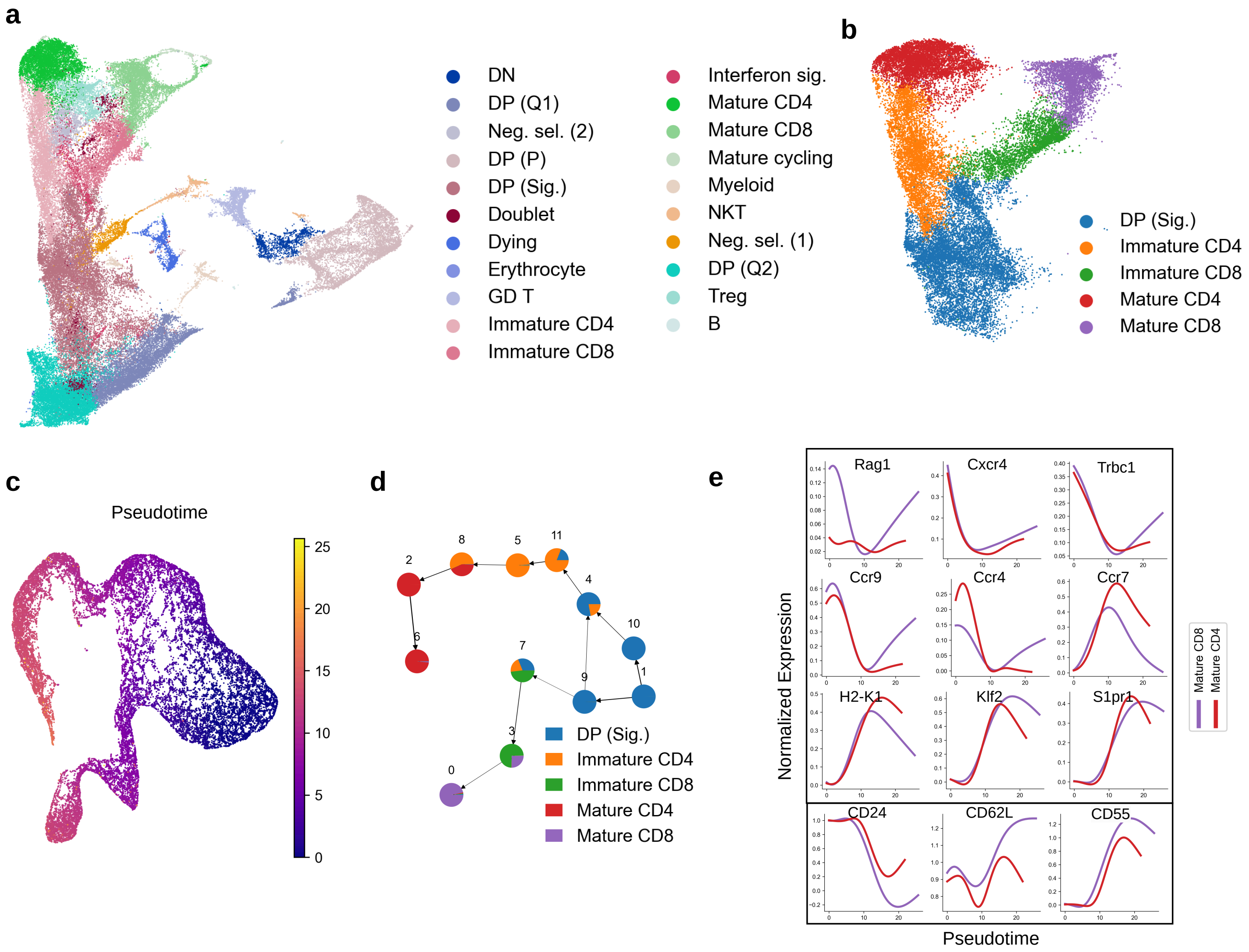


**Supplementary Figure 4**: **Application of TotalVI on a thymocyte development dataset.** (a) UMAP visualization of TotalVI embeddings for the Thymocyte development dataset, with cells colored according to cell type. (b) Subset of positive selection cells from (a) for trajectory inference. (c) Trajectory of positively selected CD4+ and CD8+ T cells inferred by MARGARET based on TotalVI-inferred embeddings. (d) UMAP plot of trajectory delineating the progression of pseudotime. (e) Gene expression trends for known marker genes for the CD4+ and CD8+ T cell lineages using the denoised expression values from TotalVI, scaled per gene.

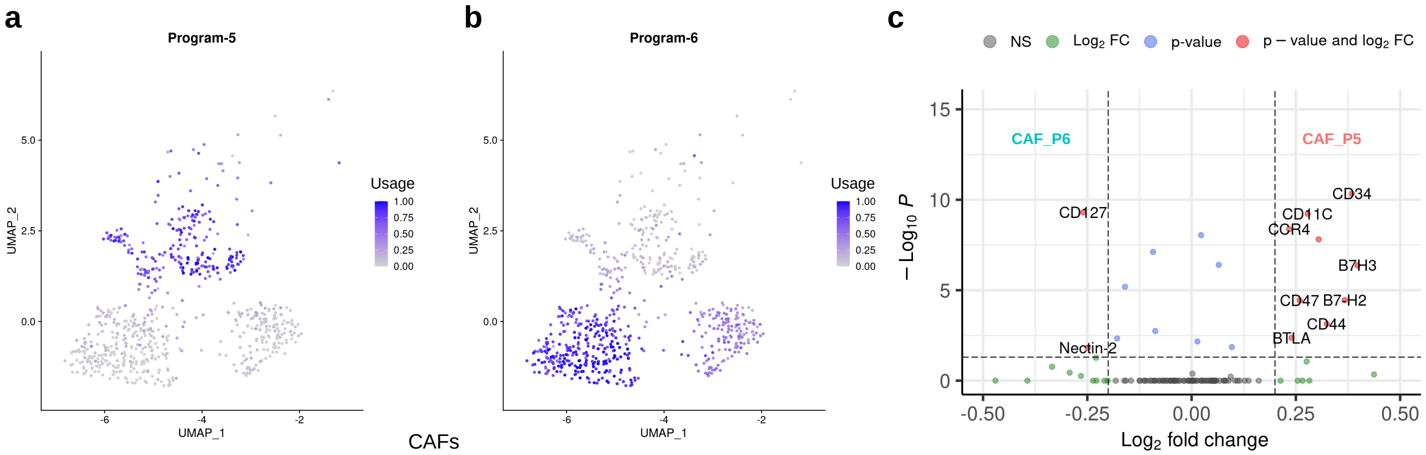


**Supplementary Figure 5**: UMAP plot of cancer associated fibroblasts (CAFs) showing the usage of (a) Program-5, and (b) Program-6 encoded in the shared latent embeddings ($\boldsymbol{z}^{s_1}$) inferred by multiHIVE for the breast cancer dataset. (c) Volcano plot of differentially expressed proteins in the CAF subpopulations (CAF_P5 and CAF_P6).

### Supplementary Tables

**Supplementary Table 1**: Pathways associated with cell type-specific gene expression programs corresponding to multiHIVE embedding $\boldsymbol{z}^{\boldsymbol{s_1}}$ for Thymocyte development dataset and their corresponding references.

| Cell type | Pathway | Reference |
|---|---|---|
| DP (Q1), DP (Q2) | TRANSLOCATION_OF_ZAP_70_TO_IMMUNOLOGICAL SYNAPSE | (1) |
| | PHOSPHORYLATION_OF_CD3_AND_TCR_ZETA_CHAINS | (2; 3) |
| DP (Sig.) | TNFA_SIGNALING_VIA_NFKB | (4; 5) |
| Mature CD4 T cells, Mature CD8 T cells | CD8_POSITIVE_ALPHA_BETA_T_CELL_ACTIVATION | (6) |
| | T_CELL_MEDIATED_CYTOTOXICITY | (7) |
| | IL2_STAT5_SIGNALING | (8; 9) |
| $\gamma\delta$ T cells | TRANSMEMBRANE_RECEPTOR_PROTEIN_TYROSINE KINASE_SIGNALING_PATHWAY | (10; 11) |

### Supplementary References


[1] Palacios, E. H. & Weiss, A. Distinct roles for syk and zap-70 during early thymocyte development. *The Journal of experimental medicine* **204**, 1703–1715 (2007).

[2] Levelt, C., Carsetti, R. & Eichmann, K. Regulation of thymocyte development through cd3. ii. expression of t cell receptor beta cd3 epsilon and maturation to the cd4+ 8+ stage are highly correlated in individual thymocytes. *The Journal of experimental medicine* **178**, 1867–1875 (1993).

[3] Brodeur, J.-F., Li, S., Damlaj, O. & Dave, V. P. Expression of fully assembled tcr–cd3 complex on double positive thymocytes: synergistic role for the prs and er retention motifs in the intra-cytoplasmic tail of cd3$\varepsilon$. *International immunology* **21**, 1317–1327 (2009).

[4] Webb, L. V., Ley, S. C. & Seddon, B. Tnf activation of nf-$\kappa$b is essential for development of single-positive thymocytes. *Journal of Experimental Medicine* **213**, 1399–1407 (2016).

[5] Liu, T., Zhang, L., Joo, D. & Sun, S.-C. Nf-$\kappa$b signaling in inflammation. *Signal transduction and targeted therapy* **2**, 1–9 (2017).

[6] Lambolez, F., Kronenberg, M. & Cheroutre, H. Thymic differentiation of tcr$\alpha\beta$+ cd8$\alpha\alpha$+ iels. *Immunological reviews* **215**, 178–188 (2007).

[7] Overgaard, N. H., Jung, J.-W., Steptoe, R. J. & Wells, J. W. Cd4+/cd8+ double-positive t cells: more than just a developmental stage? *Journal of Leucocyte Biology* **97**, 31–38 (2015).

[8] Moriggl, R. *et al.* Stat5 is required for il-2-induced cell cycle progression of peripheral t cells. *Immunity* **10**, 249–259 (1999).

[9] Mahmud, S. A., Manlove, L. S. & Farrar, M. A. Interleukin-2 and stat5 in regulatory t cell development and function. *Jak-Stat* **2**, e23154 (2013).

[10] Parker, M. E. & Ciofani, M. Regulation of $\gamma\delta$ t cell effector diversification in the thymus. *Frontiers in immunology* **11**, 42 (2020).

[11] Muro, R., Takayanagi, H. & Nitta, T. T cell receptor signaling for $\gamma\delta$t cell development. *Inflammation and Regeneration* **39**, 1–11 (2019).
